## Supplemental figures and tables for "Detecting genetic variation and base modifications together in the same single molecules of DNA and RNA at base pair resolution using a magnetic tweezer platform"

### Supplemental material

#### Figures and tables

***Supplemental Figure 1.*** *Estimation of the platform noise.*

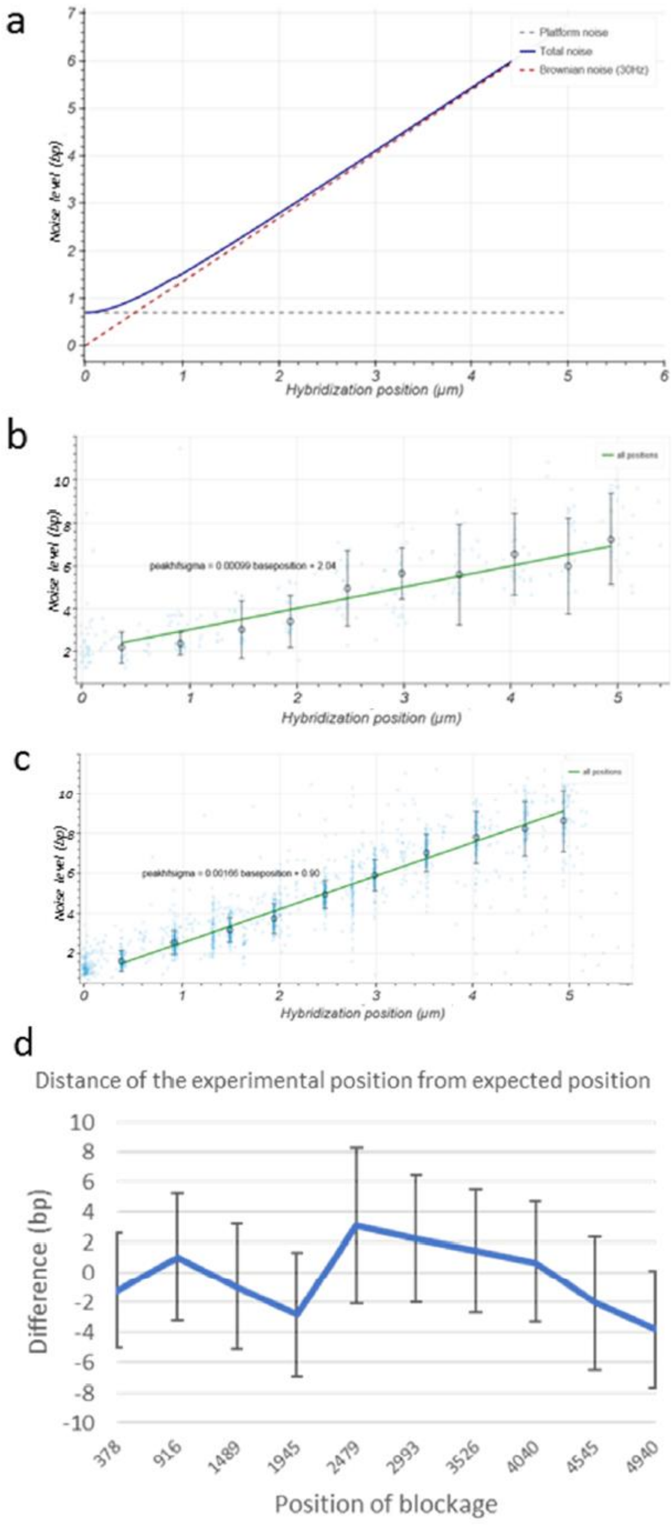

**(a)** Modelling of the measured total noise (dark blue line), which is composed of the platform noise (which is constant independently of the molecule length, black dotted line) plus the Brownian noise, which is dependent on the length of the molecule (red dotted line). A 5 kb hairpin was constructed to determine the platform noise by testing ten ten-base oligonucleotides along the length of the molecules. For each position, the noise was recorded at a frequency of 30 images per second and plotted against the position of blockage (each bleu point corresponds to a mapped blocking position). After fitting a linear regression, it is possible to determine the instrument noise (when the molecule length is 0), which is 2.04 bases for the original MT **(b)** and 0.90 bases for the new SDI platform **(c)**. **(d)** The difference in base pairs between the experimental and expected position for all the oligonucleotides along the 5 kb hairpin is plotted. For positions up to 1.5 kb from the Y-shape, the precision is less than 1 base and reached four bases at the 4940 bp position of the hairpin.

***Supplemental Figure 2.*** *Principle of the loop PCR strategy.*

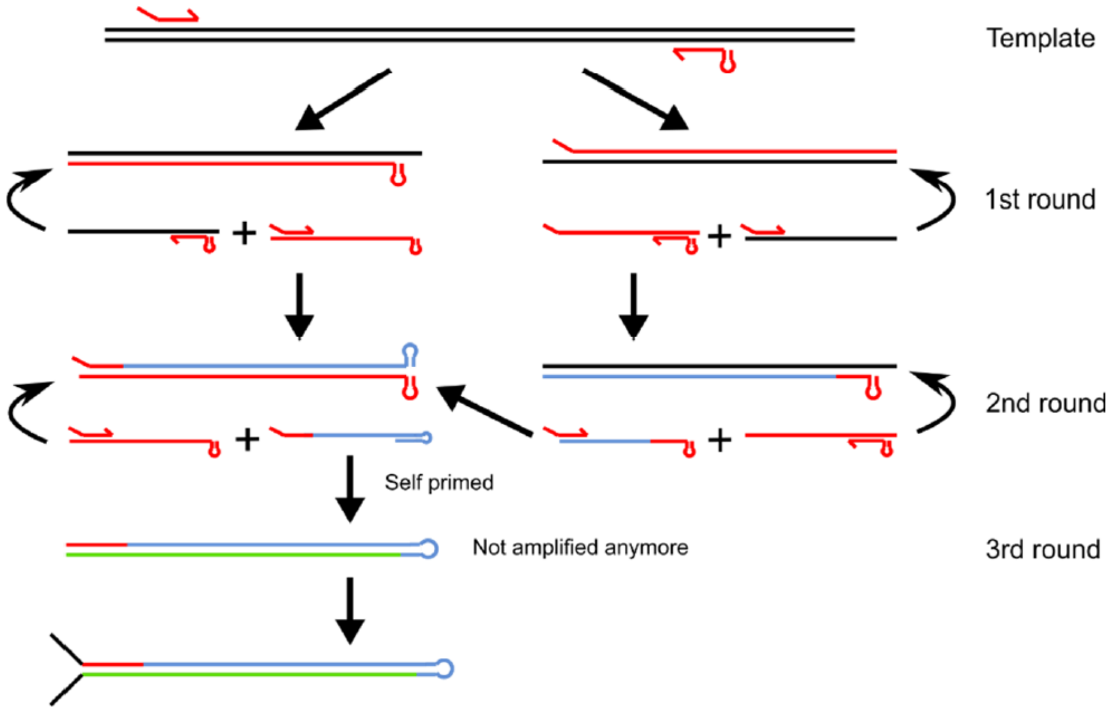


The loop PCR strategy uses a normal forward primer, but the reverse primer is designed so that it encompasses a loop. At the first cycle of the PCR, the reverse looped primer creates a DNA molecule which incorporates the loop at its 5' end, which cannot be extended. In the second cycle, when the forward primer is extended, it will copy the loop generated in the previous cycle. In the third cycle, the replicated loop, which is now at the 3' end of the transcript, will fold back on itself and self-prime. Since this newly synthesized molecule is in a hairpin form it cannot be further amplified. The remaining single-stranded cDNAs with loops at their 5' ends will act as templates to each form another 3' loop molecule, which leads to the desired double-stranded hairpin product.

***Supplemental Figure 3.*** *Quantification of CAPZB in different cellular state using different techniques.*

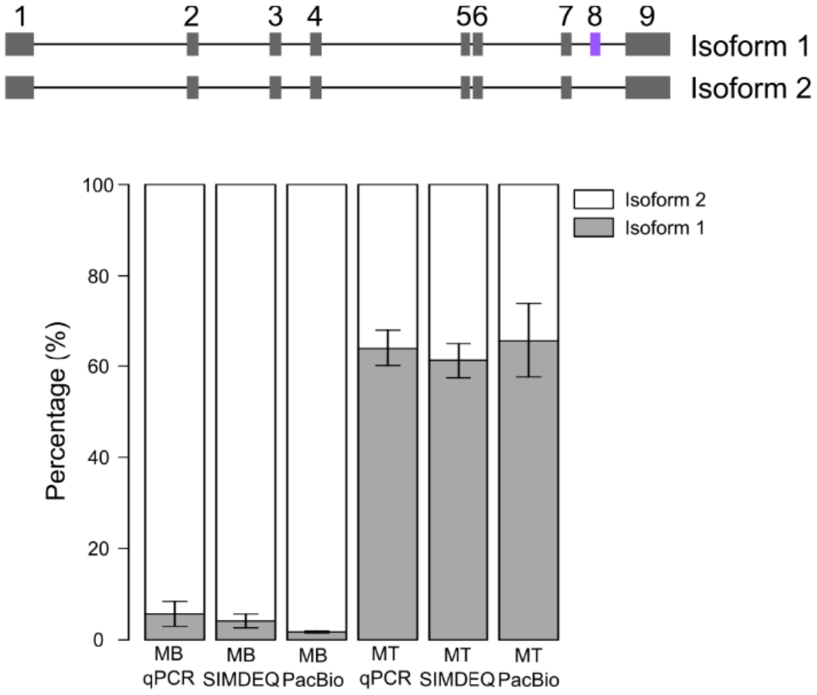


Both isoforms of CAPXB were quantified in both myoblast and myotube for the inclusion or exclusion of exon 8. Similar results were obtained independently of the technique used for quantification.

***Supplemental Figure 4.*** *Cas meganucleases can shield the end of DNA fragments from digestion with exonucleases.*

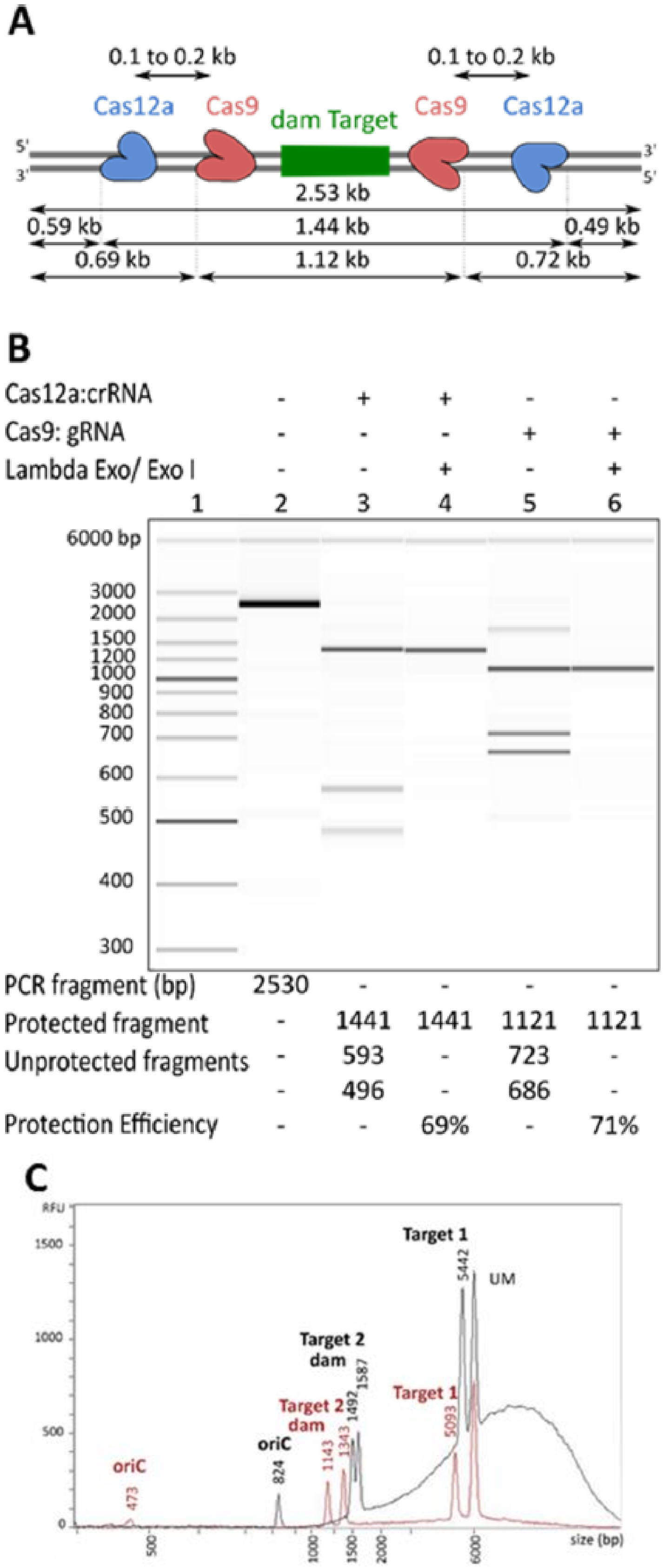

**(A)** Schematic representation of the experimental design of the protection assay. PCR fragment was generated such that the left- and right-side fragments resulting from digestion with Cas protein are of different sizes, allowing their simultaneous detection by capillary electrophoresis (schema not to scale). **(B)** The PCR product was incubated with either the Cas12a:crRNA complex (lanes 3-4) or the cas9:gRNA complex (lanes 5-6). After one hour of incubation, the reaction was supplemented with lambda Exo/ExoI nucleases (lanes 4 and 6) to digest all the DNA located outside the two Cas complexes. The expected fragments sizes and protection efficacy for each complex are listed below the Figure. **(C)** The protection of four targets from *E. coli* was achieved by incubating genomic DNA with all eight Cas12a:crRNA complexes corresponding to the four targets and after one hour, the reaction was supplemented with a mix of exonucleases (black outline). After this first protection step, 1/10th of the protected material was incubated with the eight dCas9:gRNA complexes and after one hour of incubation, exonucleases were added to the reaction tube (red outline). The peak corresponding to each target is marked as well as their estimated sizes. After the first protection step, a large quantity of undigested gDNA is present, indicated by the dome. This is eliminated with the second protection step. RFU: relative fluorescence unit. UM: Upper marker.

***Supplemental Figure 5.*** *Signature produced by the oligonucleotide CAAG on the enriched targets for human gDNA.*

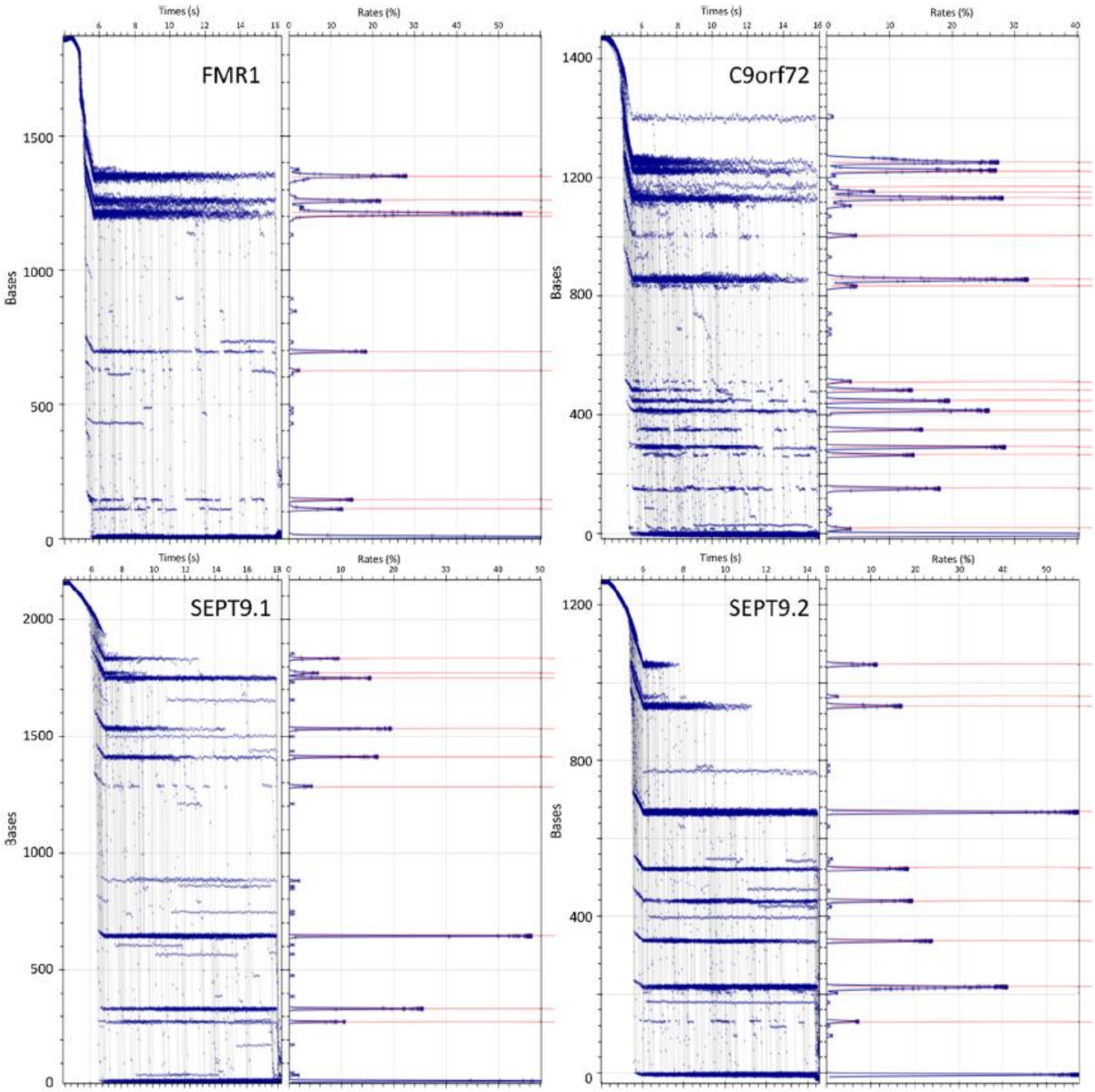


Expected and experimental positions of the CAAG oligonucleotide on the FMR1, C9orf72, SEPT9.1 and SEPT9.2 hairpin produced from the enriched fragments. Each red line represents one expected blockage. These specific patterns allowed us to identify and classify the functional hairpins as one of the four targets.

***Supplemental Figure 6.*** *Principle of the repeat analysis of FMR1 using an opening assay.*

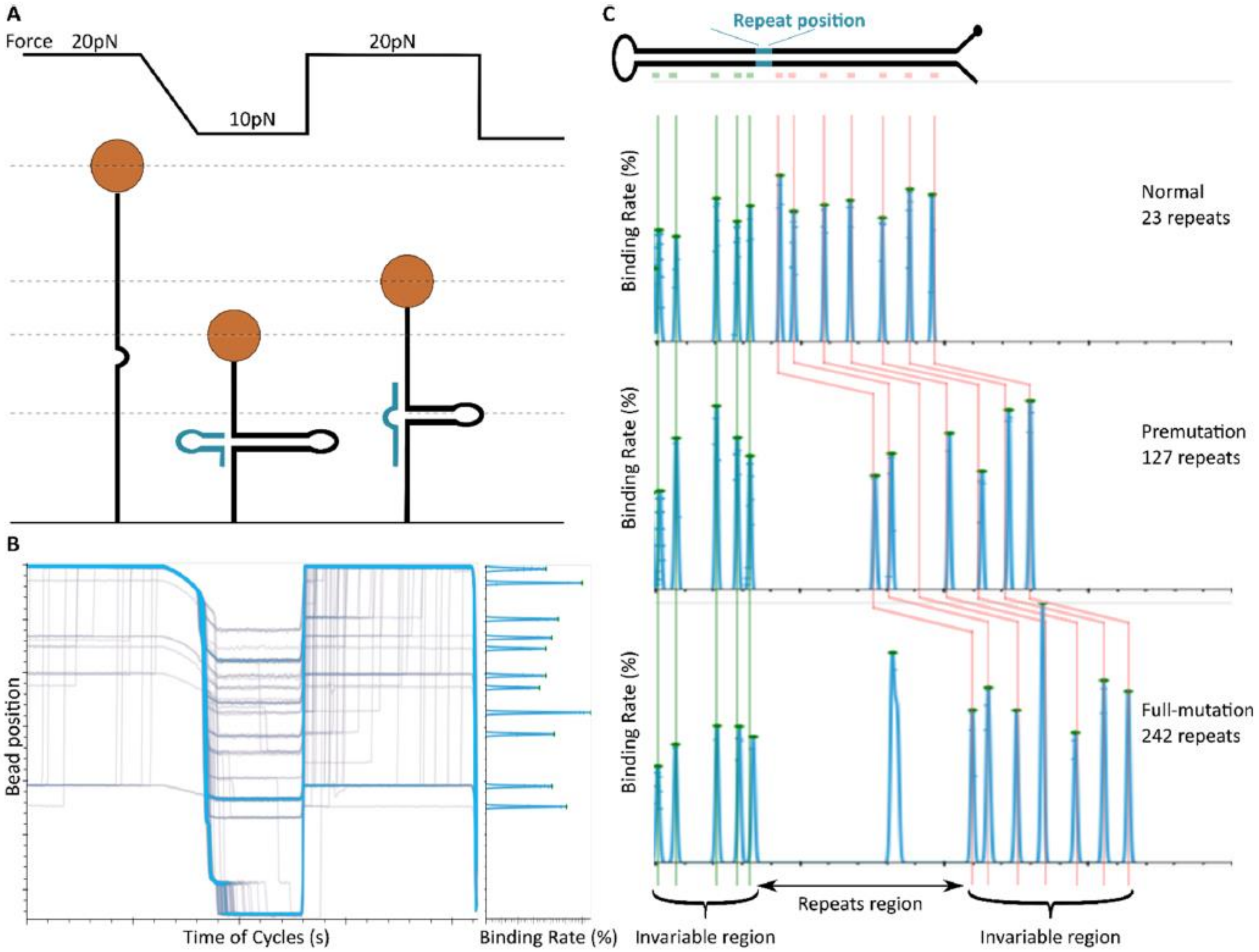


**(A)** An oligonucleotide designed to form a three-way junction is injected into the flow cell and the functional molecules are cycled between high and low forces. The oligonucleotides bind to the molecules during the phase of high force (20 pN) through a single-stranded end and cause a transient blockage when the force is reduced. When the force is increased to 20 pN, a three-way junction is formed and transiently prevents opening of the molecules. Measurement at high force prevents any secondary structures (for example, intra-molecule hairpins or G-quadruplexes) that could affect the measurement of the blocking position during the closing phase. **(B)** Typical opening and closing cycles for this assay (left part) and the corresponding histogram of the detected blocking positions (right part). **(C)** Schematic representation of the invariable positions of the reference oligonucleotides that bind on either side of the repeat region of the FMR1 hairpin. Green lines represent the theoretical binding position of the five oligonucleotides located upstream of the repeats and the red lines are the positions located downstream of the repeats. The extracted experimental peaks were fitted to these reference positions (X-axis) and used to calibrate the measurement of the repeat length. The graph represents three different molecules with either 23, 127 or 242 repeats. (The additional peak within the repeat region of the molecule with 242 repeats is due to a non-specific blockage of the molecule within the repeats.)

***Supplemental Table 1.*** *CpG and Non-CpG sites on the FMR1 locus.*

| | | | | Methylated Molecules | | | | | |
|---|---|---|---|---|---|---|---|---|---|
| | | | | WT n=11 | | Pre-Mutated n=9 | | Full-Mutated n=3 | |
| Site Number | Position within HP | Sequence | Chr Position | Nb | Ratio | Nb | Ratio | Nb | Ratio |
| 1 | 276 | CG | 147911340 | 1 | 0.09 | 0 | 0.00 | 0 | 0.00 |
| 2 | 300 | CG | 147911364 | 3 | 0.27 | 3 | 0.33 | 0 | 0.00 |
| 3 | 306 | CG | 147911370 | 1 | 0.09 | 3 | 0.33 | 0 | 0.00 |
| 4 | 321 | CT | 147911385 | 0 | 0.00 | 1 | 0.11 | 0 | 0.00 |
| 5 | 326 | CG | 147911390 | 2 | 0.18 | 1 | 0.11 | 0 | 0.00 |
| 6 | 346 | CA | 147911410 | 0 | 0.00 | 1 | 0.11 | 0 | 0.00 |
| 7 | 388 | CT | 147911452 | 0 | 0.00 | 2 | 0.22 | 0 | 0.00 |
| 8 | 397 | CA | 147911461 | 0 | 0.00 | 2 | 0.22 | 0 | 0.00 |
| 9 | 409 | CG | 147911473 | 3 | 0.27 | 2 | 0.22 | 1 | 0.33 |
| 10 | 428 | CG | 147911492 | 5 | 0.45 | 2 | 0.22 | 0 | 0.00 |
| 11 | 440 | CT | 147911504 | 0 | 0.00 | 1 | 0.11 | 0 | 0.00 |
| 12 | 443 | CG | 147911507 | 3 | 0.27 | 2 | 0.22 | 0 | 0.00 |
| 13 | 448 | CT | 147911512 | 0 | 0.00 | 1 | 0.11 | 0 | 0.00 |
| 14 | 456 | CG | 147911520 | 4 | 0.36 | 3 | 0.33 | 0 | 0.00 |
| 15 | 510 | CG | 147911574 | 4 | 0.36 | 4 | 0.44 | 0 | 0.00 |
| 16 | 512 | CG | 147911576 | 4 | 0.36 | 4 | 0.44 | 0 | 0.00 |
| 17 | 529 | CG | 147911593 | 3 | 0.27 | 2 | 0.22 | 1 | 0.33 |
| 18 | 543 | CG | 147911607 | 3 | 0.27 | 2 | 0.22 | 0 | 0.00 |
| 19 | 563 | CG | 147911627 | 6 | 0.55 | 2 | 0.22 | 0 | 0.00 |
| 20 | 593 | CG | 147911657 | 6 | 0.55 | 3 | 0.33 | 0 | 0.00 |
| 21 | 605 | CG | 147911669 | 5 | 0.45 | 2 | 0.22 | 0 | 0.00 |
| 22 | 607 | CG | 147911671 | 5 | 0.45 | 2 | 0.22 | 0 | 0.00 |
| 23 | 617 | CG | 147911681 | 4 | 0.36 | 3 | 0.33 | 0 | 0.00 |
| 24 | 625 | CA | 147911689 | 1 | 0.09 | 1 | 0.11 | 0 | 0.00 |
| 25 | 628 | CG | 147911692 | 5 | 0.45 | 5 | 0.56 | 0 | 0.00 |
| 26 | 639 | CT | 147911703 | 0 | 0.00 | 1 | 0.11 | 0 | 0.00 |
| 27 | 644 | CG | 147911708 | 5 | 0.45 | 4 | 0.44 | 0 | 0.00 |
| 28 | 660 | CG | 147911724 | 4 | 0.36 | 3 | 0.33 | 0 | 0.00 |
| 29 | 662 | CG | 147911726 | 3 | 0.27 | 3 | 0.33 | 0 | 0.00 |
| 30 | 666 | CG | 147911730 | 4 | 0.36 | 2 | 0.22 | 0 | 0.00 |
| 31 | 670 | CG | 147911734 | 2 | 0.18 | 3 | 0.33 | 0 | 0.00 |
| 32 | 675 | CG | 147911739 | 4 | 0.36 | 3 | 0.33 | 0 | 0.00 |
| 33 | 677 | CG | 147911741 | 4 | 0.36 | 3 | 0.33 | 0 | 0.00 |
| 34 | 679 | CG | 147911743 | 4 | 0.36 | 3 | 0.33 | 0 | 0.00 |
| 35 | 690 | CG | 147911754 | 5 | 0.45 | 2 | 0.22 | 0 | 0.00 |
| 36 | 695 | CG | 147911759 | 2 | 0.18 | 2 | 0.22 | 0 | 0.00 |
| 37 | 703 | CG | 147911767 | 4 | 0.36 | 2 | 0.22 | 0 | 0.00 |
| 38 | 707 | CG | 147911771 | 3 | 0.27 | 3 | 0.33 | 0 | 0.00 |
| 39 | 714 | CA | 147911778 | 1 | 0.09 | 2 | 0.22 | 0 | 0.00 |
| 40 | 720 | CG | 147911784 | 2 | 0.18 | 2 | 0.22 | 0 | 0.00 |
| 41 | 722 | CG | 147911786 | 1 | 0.09 | 2 | 0.22 | 0 | 0.00 |
| 42 | 728 | CG | 147911792 | 2 | 0.18 | 2 | 0.22 | 0 | 0.00 |
| 43 | 730 | CG | 147911794 | 2 | 0.18 | 0 | 0.00 | 0 | 0.00 |
| 44 | 732 | CG | 147911796 | 2 | 0.18 | 1 | 0.11 | 0 | 0.00 |
| 45 | 746 | CA | 147911810 | 0 | 0.00 | 3 | 0.33 | 0 | 0.00 |
| 46 | 750 | CT | 147911814 | 5 | 0.45 | 1 | 0.11 | 0 | 0.00 |
| 47 | 760 | CG | 147911824 | 5 | 0.45 | 4 | 0.44 | 0 | 0.00 |
| 48 | 766 | CG | 147911830 | 5 | 0.45 | 5 | 0.56 | 0 | 0.00 |
| 49 | 780 | CG | 147911844 | 3 | 0.27 | 3 | 0.33 | 0 | 0.00 |
| 50 | 782 | CG | 147911846 | 3 | 0.27 | 3 | 0.33 | 0 | 0.00 |
| 51 | 798 | CG | 147911862 | 3 | 0.27 | 4 | 0.44 | 0 | 0.00 |
| 52 | 807 | CG | 147911871 | 3 | 0.27 | 2 | 0.22 | 2 | 0.67 |
| 53 | 812 | CG | 147911876 | 4 | 0.36 | 2 | 0.22 | 2 | 0.67 |
| 54 | 815 | CA | 147911879 | 1 | 0.09 | 1 | 0.11 | 0 | 0.00 |
| 55 | 820 | CA | 147911884 | 3 | 0.27 | 2 | 0.22 | 0 | 0.00 |
| 56 | 829 | CA | 147911893 | 4 | 0.36 | 3 | 0.33 | 0 | 0.00 |
| 57 | 833 | CG | 147911897 | 3 | 0.27 | 2 | 0.22 | 0 | 0.00 |
| 58 | 838 | CG | 147911902 | 4 | 0.36 | 3 | 0.33 | 0 | 0.00 |
| 59 | 843 | CG | 147911907 | 3 | 0.27 | 4 | 0.44 | 0 | 0.00 |
| 60 | 851 | CG | 147911915 | 4 | 0.36 | 4 | 0.44 | 0 | 0.00 |
| 61 | 856 | CA | 147911920 | 2 | 0.18 | 4 | 0.44 | 0 | 0.00 |
| 62 | 865 | CG | 147911929 | 3 | 0.27 | 5 | 0.56 | 0 | 0.00 |
| 63 | 875 | CG | 147911939 | 6 | 0.55 | 6 | 0.67 | 0 | 0.00 |
| 64 | 880 | CG | 147911944 | 5 | 0.45 | 6 | 0.67 | 0 | 0.00 |
| 65 | 892 | CG | 147911956 | 4 | 0.36 | 5 | 0.56 | 0 | 0.00 |
| 66 | 899 | CT | 147911963 | 0 | 0.00 | 4 | 0.44 | 0 | 0.00 |
| 67 | 903 | CG | 147911967 | 4 | 0.36 | 5 | 0.56 | 0 | 0.00 |
| 68 | 913 | CG | 147911977 | 5 | 0.45 | 7 | 0.78 | 0 | 0.00 |
| 69 | 917 | CG | 147911981 | 4 | 0.36 | 4 | 0.44 | 0 | 0.00 |
| 70 | 920 | CG | 147911984 | 3 | 0.27 | 2 | 0.22 | 0 | 0.00 |
| 71 | 925 | CG | 147911989 | 4 | 0.36 | 5 | 0.56 | 0 | 0.00 |
| 72 | 928 | CG | 147911992 | 4 | 0.36 | 5 | 0.56 | 0 | 0.00 |
| 73 | 931 | CG | 147911995 | 5 | 0.45 | 3 | 0.33 | 0 | 0.00 |
| 74 | 935 | CG | 147911999 | 1 | 0.09 | 5 | 0.56 | 0 | 0.00 |
| 75 | 937 | CG | 147912001 | 0 | 0.00 | 2 | 0.22 | 0 | 0.00 |
| 76 | 941 | CG | 147912005 | 3 | 0.27 | 4 | 0.44 | 0 | 0.00 |
| 77 | 944 | CG | 147912008 | 3 | 0.27 | 3 | 0.33 | 0 | 0.00 |
| 78 | 947 | CG | 147912011 | 3 | 0.27 | 3 | 0.33 | 0 | 0.00 |
| 79 | 953 | CG | 147912017 | 4 | 0.36 | 5 | 0.56 | 0 | 0.00 |
| 80 | 957 | CT | 147912021 | 2 | 0.18 | 1 | 0.11 | 0 | 0.00 |
| 81 | 959 | CG | 147912023 | 4 | 0.36 | 3 | 0.33 | 0 | 0.00 |
| 82 | 962 | CG | 147912026 | 3 | 0.27 | 5 | 0.56 | 0 | 0.00 |
| 83 | 963 | CT | 147912027 | 3 | 0.27 | 2 | 0.22 | 0 | 0.00 |
| 84 | 968 | CA | 147912032 | 3 | 0.27 | 3 | 0.33 | 0 | 0.00 |
| 85 | 975 | CG | 147912039 | 4 | 0.36 | 3 | 0.33 | 0 | 0.00 |
| 86 | 979 | CG | 147912043 | 3 | 0.27 | 5 | 0.56 | 0 | 0.00 |
| 87 | 985 | CG | 147912049 | 4 | 0.36 | 5 | 0.56 | 0 | 0.00 |

List of all the positions and the rate of methylation across the molecule of the DNA sample NA06896.
